# Formation of tRNA wobble inosine in humans is perturbed by a millennia-old mutation linked to intellectual disability

**DOI:** 10.1101/277079

**Authors:** Jillian Ramos, Lu Han, Yan Li, Fowzan S. Alkuraya, Eric M. Phizicky, Dragony Fu

**Affiliations:** Department of Biology, University of Rochester, King Faisal Specialist Hospital and Research Center, Riyadh, Saudi Arabia; Center for RNA Biology, King Faisal Specialist Hospital and Research Center, Riyadh, Saudi Arabia; Department of Biochemistry and Biophysics, University of Rochester Medical Center, King Faisal Specialist Hospital and Research Center, Riyadh, Saudi Arabia; Department of Genetics, King Faisal Specialist Hospital and Research Center, Riyadh, Saudi Arabia; Department of Anatomy and Cell Biology, College of Medicine, Alfaisal University, Riyadh, Saudi Arabia

## Abstract

The formation of inosine at the wobble position of eukaryotic tRNAs is an essential modification catalyzed by the ADAT2/ADAT3 complex. In humans, a valine to methionine mutation (V144M) in ADAT3 that originated ∼1,600 years ago is the most common cause of autosomal-recessive intellectual disability (ID) in Arabia. Here, we show that ADAT3-V144M exhibits perturbations in subcellular localization and has increased propensity to form aggregates associated with cytoplasmic chaperonins. While ADAT2 co-expression can suppress the aggregation of ADAT3-V144M, the ADAT2/3 complexes assembled with ADAT3-V144M exhibit defects in adenosine deaminase activity. Moreover, extracts from cell lines derived from ID-affected individuals expressing only ADAT3-V144M display a reduction in tRNA deaminase activity. Notably, we find that the same cell lines from ID-affected individuals exhibit decreased wobble inosine in certain tRNAs. These results identify a role for ADAT2-dependent localization and folding of ADAT3 in wobble inosine modification that is crucial for the developing human brain.

## Introduction

The hydrolytic deamination of adenosine (A) to inosine (I) at the wobble position of tRNA is an essential post-transcriptional tRNA modification in bacteria and eukaryotes (reviewed in (1–3). Since inosine can pair with U, C or A, a single tRNA isoacceptor containing the inosine modification at the wobble anticodon position can recognize up to three different codons containing a different nucleotide base at the third position. Thus, the degeneracy provided by the wobble inosine modification is necessary for the translation of C or A-ending codons in organisms that lack a cognate G or U_34_-containing anticodon tRNA isoacceptor by expanding the reading capacity of tRNA isoacceptors (4). Moreover, it has been shown that highly translated genes in eukaryotic organisms, including humans, are correlated with an enrichment in wobble inosine tRNA-dependent codons, suggesting a critical role for tRNA inosine modification in maintaining proper levels of protein expression (5,6).

In *Escherichia coli*, A to I conversion at the wobble position is present in a single tRNA (tRNA-Arg-ACG); and is catalyzed by the homodimeric complex TadA adenosine deaminase (7). In the yeast *Saccharomyces cerevisiae*, wobble inosine modification occurs in seven different tRNAs and is catalyzed by a heterodimeric enzyme complex consisting of the Tad2p and Tad3p subunits (8,9). Tad2p is the catalytic subunit and contains a prototypical deaminase motif homologous to cytidine/deoxycytidine deaminases, including a conserved glutamic acid residue within the active site that is necessary for proton shuttling in the hydrolytic deamination reaction (10,11). Tad3p also contains a canonical deaminase motif but lacks the conserved catalytic glutamate in the active site. However, Tad2p is inactive without Tad3p, indicating that formation of a heterodimeric Tad2p/Tad3p complex is required for adenosine deaminase activity (8). Functional homologs of *S. cerevisiae* Tad2p and Tad3p have been identified in all eukaryotes to date, including the human homologs ADAT2 and ADAT3 (12–15).

Exome sequencing and autozygosity mapping have identified a single c.382G∼A mutation in the human *ADAT3* gene that is causative for autosomal-recessive intellectual disability (ID) in multiple families of Saudi Arabian descent (16,17). All reported individuals homozygous for the V144M mutation exhibit cognitive deficits indicative of a neurodevelopmental disorder, with the majority displaying strabismus and growth delay. Additional clinical features of individuals homozygous for the ADAT3-V144M mutation include microcephaly, epilepsy, and occasional brain abnormalities such as white matter atrophy and arachnoid cysts. Subsequent large-scale sequencing has identified this ancient founder mutation to be one of the most common causes of autosomal recessive intellectual disability in patients from Saudi Arabia, with a carrier frequency of ∼1% (18–20). However, the mechanistic cause of ADAT3-associated pathogenesis remains unclear.

Based upon the longer of two mRNA transcripts encoded by the human *ADAT3* gene, the ID-causing G∼A transition results in a valine to methionine missense mutation at residue 144 (V144M) of the encoded ADAT3 protein. The mutated valine residue is conserved from yeast to humans and is predicted to perturb the surface structure of the ADAT3 protein (16). However, it is unknown how the V144M mutation affects ADAT3 function and whether this would affect tRNA inosine modification levels in ID-affected individuals that are homozygous for the autosomal-recessive mutation. This would be important to test given the increasing awareness of tRNA modification in the etiology of other forms of Mendelian ID (21–30).

Here, we use subcellular localization studies combined with proteomics to find that ADAT3-V144M exhibits aberrant aggregation into cytoplasmic foci accompanied by targeting by chaperonin complexes HSP60 and TRiC/CCT. Interestingly, the aggregation phenotype of ADAT3-V144M along with its association with chaperonins can be suppressed by co-expression with ADAT2. Moreover, we find that purified ADAT2/3 complexes assembled with ADAT3-V144M display severe defects in adenosine deaminase activity on a known tRNA substrate. Most strikingly, we find that cells isolated from ID-affected individuals homozygous for the ADAT3-V144M mutation contain diminished levels of wobble inosine in several tRNA isoacceptors and extracts from these cells exhibit a drastic decrease in adenosine deaminase activity. Altogether, these results uncover a molecular basis for ADAT3-associated neurodevelopmental disorders in the form of diminished inosine modifications at the wobble position of tRNA caused by aberrant localization of the ADAT2/3 adenosine deaminase complex combined with defects in enzymatic activity.

## RESULTS

### ADAT3-V144M displays aberrant subcellular localization and increased susceptibility to form cytoplasmic aggregates

While the ID-associated V144M mutation is predicted to affect protein structure, it is unknown if and how the V144M mutation affects the function of ADAT3 in tRNA modification. Since wobble inosine modification has been proposed to occur at multiple stages of tRNA maturation in the nucleus and cytoplasm of eukaryotes (14,15,31), we first monitored whether the subcellular localization of ADAT3 was perturbed by the V144M mutation. The localization of ADAT3 was analyzed by microscopy of HeLa human cervical carcinoma cells transiently expressing ADAT3 fusion proteins with green-fluorescent protein at the amino-terminus (GFP-ADAT3). Whereas GFP alone displayed uniform accumulation in both the cytoplasm and nucleus of Hela cells (Figure 1A, GFP), the majority of cells expressing GFP-ADAT3-wildtype (WT) exhibited diffuse cytoplasmic localization outlining the nucleus with only a small percentage of transfected cells exhibiting GFP-ADAT3 signal in the nucleus (Figure 1A, B, GFP-ADAT3-WT). The absence of nuclear localization for GFP-ADAT3-WT is likely due to the limiting amounts of endogenous ADAT2 subunit that is required for nuclear import of ADAT3 (15). By contrast, the ADAT3-V144M variant exhibited a distinct localization pattern with distribution in both the cytoplasm and nucleus rather than the primarily cytoplasmic localization of ADAT3-WT (Figure 1A, B). In addition to aberrant nuclear localization, we detected an increased population of GFP-positive cells that exhibited discrete, cytoplasmic foci when transfected with the ADAT3-V144M variant (Figure 1A, C; GFP-ADAT3-V144M, arrowheads). We also found that carboxy-terminal GFP-tagged ADAT3 exhibited the same aberrant nucleocytoplasmic localization pattern and increased formation of cytoplasmic foci observed with N-terminal GFP-ADAT3 (Supplemental Figure 1). In addition, we found that the steady-state levels of GFP-ADAT3-V144M variant was lower than ADAT3-WT (Figure 1D, Supplemental Figure 1), showing that the aberrant subcellular localization pattern of ADAT3-V144M was not simply due to greater expression.

**Figure 1.**
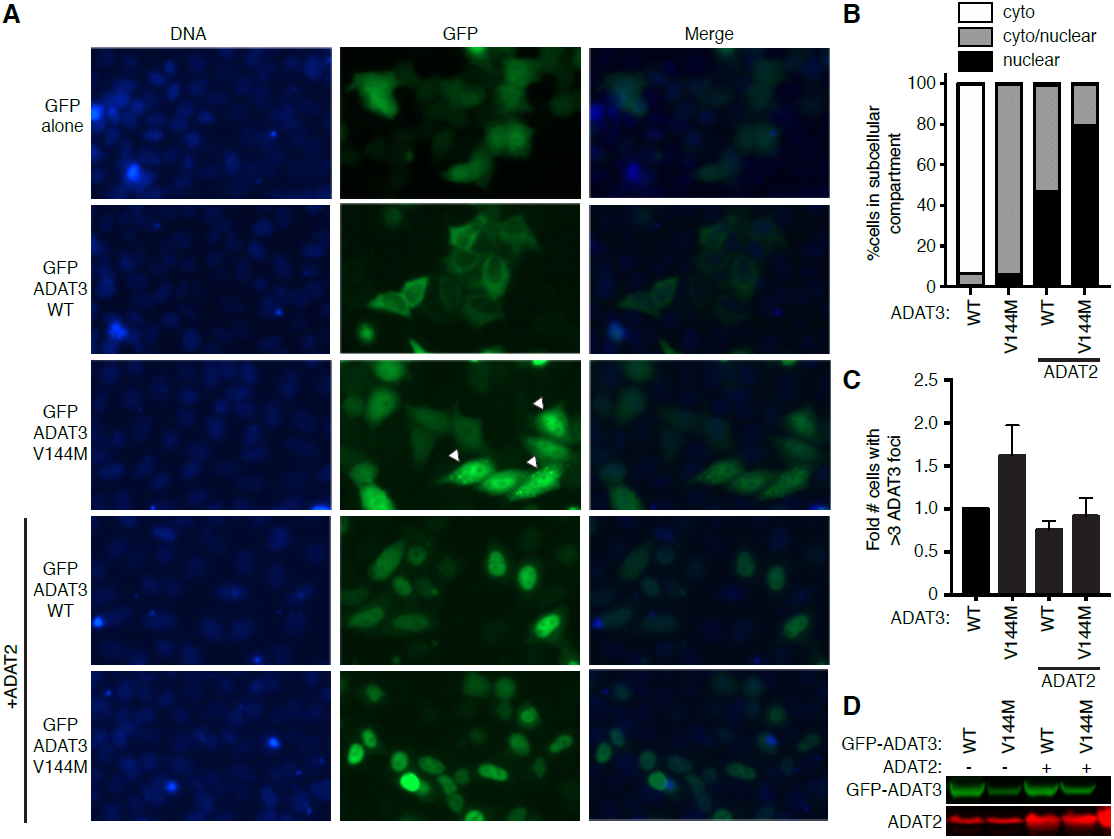
ADAT3-V144M displays aberrant nucleocytoplasmic localization and increased susceptibility to form cytoplasmic aggregates. (A) Fluorescence microscopy images of GFP alone, GFP-tagged ADAT3-WT and V144M expressed in HeLa cervical carcinoma cells. Nuclear DNA was stained with Hoechst with merge on right column. Arrowheads represent cells with ∼3 cytoplasmic foci of GFP-ADAT3. (B) Fraction of cells exhibiting GFP-ADAT3 that was either primarily cytoplasmic, similarly distributed between the cytoplasmic and nucleus, or primarily nuclear. (C) Fold number of cells that exhibited greater than three cytoplasmic foci of GFP-ADAT3. The fold amount was expressed relative to ADAT3-WT without ADAT2 co-expression where 7% of cells exhibited greater than three cytoplasmic foci. (B) and (C) were repeated 3x with a minimum of 580 cells counted per experiment. (D) Immunoblot of GFP-ADAT3 expression without or with ADAT2 co-expression.

In contrast to the cytoplasmic localization of transiently expressed GFP-ADAT3-WT alone, the co-expression of ADAT2 with ADAT3-WT led to GFP-ADAT3-WT being localized to the nucleus with only a minor proportion of signal remaining in the cytoplasm (Figure 1A, B; GFP-ADAT3-WT + ADAT2), consistent with the observation that ADAT2 dimerization with ADAT3 is required for nuclear import of the ADAT2/3 complex (15). Similarly, we found that co-expression of ADAT2 with ADAT3-V144M could also induce the translocation of GFP-ADAT3-V144M into the nucleus (Figure 1A, B; GFP-ADAT3-V144M + ADAT2). The ability of ADAT2 co-expression to induce the translocation of GFP-ADAT3-V144M into the nucleus indicates that ADAT3-V144M can still interact with ADAT2. However, while a slight diffuse signal of GFP-ADAT3-WT remained in the cytoplasm even with ADAT2 co-expression, the ADAT3-V144M variant displayed much greater nuclear accumulation in the majority of cells (Figure 1B). Remarkably, co-expression of ADAT2 with the ADAT3-V144M variant also reduced the percentage of cells with cytoplasmic GFP-ADAT3 foci to nearly wildtype levels (Figure 1C). These results uncover an aberrant subcellular localization pattern for ADAT3-V144M characterized by perturbed nucleocytoplasmic distribution and increased formation of cytoplasmic foci that can be suppressed by co-expression with ADAT2. The formation of aberrant cytoplasmic aggregates exhibited by ADAT3-V144M in the absence of ADAT2 co-expression suggests that the V144M mutation causes increased propensity for protein misfolding and aggregation which can be ameliorated by interaction with the ADAT2 subunit and import into the nucleus.

### ADAT3-V144M maintains interactions with ADAT2 but exhibits increased propensity to homo-oligomerize

The perturbed subcellular localization of ADAT3-V144M along with its accumulation in cytoplasmic foci suggests that the V144M mutation could be altering the structure of ADAT3 leading to misfolding and aggregation. Furthermore, the reduction in ADAT3-V144M cytoplasmic aggregates by ADAT2 co-expression suggests that assembly of ADAT3 with ADAT2 is critical for preventing ADAT3 self-association, especially in the context of the ADAT3-V144M variant. To analyze the interaction between ADAT3-V144M and ADAT2, we co-expressed GFP-tagged ADAT3-WT or V144M with ADAT2 tagged with the twin Strep-tag (32) in 293T human embryonic kidney cells. The Strep-tag allows for one-step affinity purification of Strep-tagged proteins on streptactin resin under native conditions followed by gentle elution with biotin to preserve any protein-protein interactions. After purification of Strep-ADAT2 on streptactin resin, we found that comparable levels of GFP-ADAT3-WT and GFP-ADAT3-V144M were interacting with ADAT2 (Figure 2A). These results indicate that ADAT3-V144M interacts with similar efficiency as ADAT3-WT with ADAT2, consistent with the ability of ADAT2 to induce the translocation of either ADAT3-WT or V144M into the nucleus (Figure 1A).

**Figure 2.**
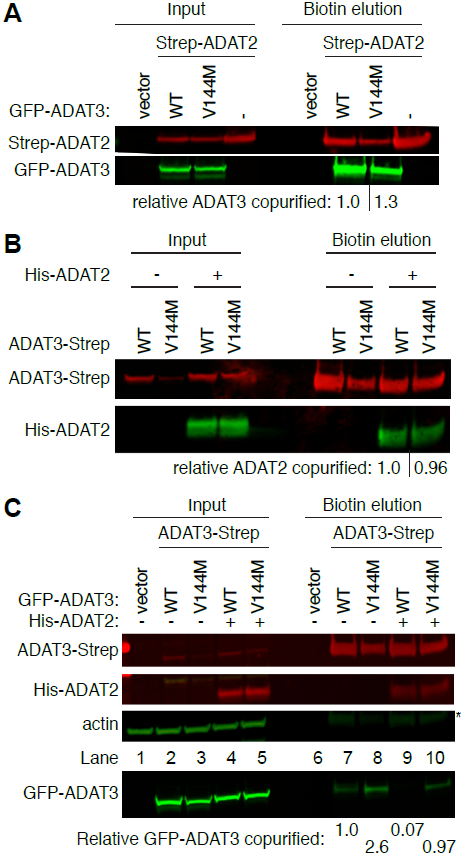
ADAT3-V144M maintains interaction with ADAT2 but displays increased propensity to self-associate. (A) Similar levels of either ADAT3-WT or ADAT3-V144M copurify with ADAT2. Immunoblot for the indicated proteins from input (5%) or Streptactin affinity purifications (10%) from 293T cells transfected to express the Strep-tag alone (vector) or Strep-tagged ADAT2 with either GFP-ADAT3-WT or V144M. The relative ADAT3 copurified represents the ratio of GFP-ADAT3 signal present in the eluted fraction normalized to Strep-ADAT2 signal relative to ADAT3-WT. (B) ADAT3-WT or ADAT3-V144M copurify with similar levels of ADAT2. Immunoblot for the indicated proteins from input (5%) or Streptactin affinity purifications (20%) from 293T cells transfected to express ADAT3-Strep-WT or ADAT3-Strep-V144M without or with His-ADAT2. The relative ADAT2 copurified represents the ratio of His-ADAT2 signal present in the eluted fraction normalized to ADAT3-Strep signal relative to ADAT3-WT. (C) Increased self-association of ADAT3-V144M. Immunoblot for the indicated proteins from input (5%) or Streptactin affinity purifications (20%) from 293T cells transfected to express ADAT3-Strep-WT or V144M with either GFP-ADAT3-WT or V144M in the absence or presence of ADAT2 coexpression. (*) represents ADAT3-Strep signal from previous probing. The % ADAT3 copurified represents the ratio of GFP-ADAT3 signal present in the eluted fraction normalized to Strep-ADAT3 signal relative to ADAT3-WT. (A-C) were repeated three times with comparable results.

To further analyze the interaction between ADAT2 and ADAT3-V144M, we used a reciprocal approach in which we purified ADAT3 and examined the amount of co-purifying ADAT2. For these assays, we transiently expressed either ADAT3-WT or V144M fused to a carboxy-terminal Strep-tag for purification and elution of ADAT2/3. Since ADAT2 levels have been shown to be limiting for the formation of ADAT2/3 complexes in human cells (15), we co-expressed His-tagged ADAT2 with either ADAT3-Strep-WT or ADAT3-Strep-V144M to facilitate detection of any associated ADAT2. After purification on streptactin resin, bound ADAT3 complexes were eluted with biotin and analyzed by immunoblotting. Using this approach, we detected the co-purification of His-ADAT2 with either ADAT3-WT or ADAT3-V144M, consistent with the assembly of an expressed ADAT2/3 complex from the expressed proteins (Figure 2B). We detected a comparable level of His-ADAT2 that copurified with ADAT3-WT or V144M, corroborating our finding that the V144M mutation does not dramatically affect the efficiency of the interaction between ADAT2 and ADAT3.

While the association between ADAT2 and ADAT3 by co-IP analysis does not appear to be affected by the V144M mutation, the aberrant subcellular localization into cytoplasmic foci suggests that ADAT3 could have defects in folding that increases its tendency to oliogomerize and aggregate. To test for self-oligomerization, we expressed either WT or V144M versions of GFP-ADAT3 simultaneously with WT or V144M forms of ADAT3-Strep (Figure 2C, lanes 2-5) (32). We detected a low level of GFP-ADAT3-WT copurifying with ADAT3-strep-WT suggesting that ADAT3 could already be susceptible to aggregation even in the wildtype state (Figure 2C, lane 7). Notably, we found that purification of ADAT3-strep-V144M led to an increase in the amount of copurifying GFP-ADAT3-V144M compared to ADAT3-WT with itself (Figure 2C, compare lanes 7 and 8). Moreover, we found that co-expression of His-ADAT2 could suppress the self-oligomerization of ADAT3-WT while partially reducing the self-association of the ADAT3-V144M variant (Figure 2C, lanes 9 and 10). The reduction of ADAT3 self-oligomerization by ADAT2 co-expression is consistent with our observation that ADAT3-V144M foci can be suppressed by ADAT2 co-expression (Figure 1A). We also note that purification of either ADAT3-WT or V144M led to similar levels of copurifying ADAT2 (Figure 2C, ADAT2, lanes 9 and 10), consistent with our findings above that ADAT3-V144M maintains interactions with ADAT2. Altogether, these results provide further evidence that the V144M mutation increases the propensity of ADAT3 to misfold and aggregate if not properly assembled with ADAT2. The ability of ADAT2 to prevent self-association of either ADAT3-WT or V144M uncovers a critical role for proper stoichiometric levels of ADAT2 and ADAT3 to promote proper folding and nuclear import of ADAT3.

### ADAT3-V144M is targeted by the cytoplasmic HSP60 and TRiC chaperonin complexes

The perturbed subcellular localization of ADAT3-V144M combined with its increased propensity to aggregate prompted us to investigate whether ADAT3-V144M also displayed interactions with other cellular proteins. For proteomic analyses, we expressed either the WT or V144M versions of ADAT3 as fusion proteins with the FLAG epitope tag in 293T human embryonic kidney cells in order to facilitate the expression and purification of ADAT3 complexes. Following immunoprecipitation (IP), the purified samples were analyzed by SDS-PAGE and silver stain to identify ADAT3-interacting proteins. While no observable bands were found in a control purification from cells transfected with vector alone, we could detect the purification of FLAG-ADAT3-WT or V144M (Figure 3A, arrowhead) along with an additional band at ∼60 kDa specifically enriched with the ADAT3-V144M purification (Figure 3A, arrow). Proteomic analysis of the entire eluted samples from control and ADAT3 purifications by liquid chromatography-mass spectrometry (LC-MS) validated the successful recovery of ADAT3-WT or ADAT3-V144M from cellular extracts (Figure 3B, Supplemental Table 1). Notably, LC-MS analysis also revealed the copurification of heat shock protein 60 (HSP60) and all eight subunits of the TCP-1 Ring Complex (TRiC, also known CCT) with ADAT3-V144M but not ADAT3-WT (Figure 3B, Supplemental Table 1). HSP60 and TRiC subunits were identified among the top 20 scoring matches in the ADAT3-V144M purification. The HSP60 protein forms a homo-oligomeric chaperonin complex consisting of a double-heptameric ring that associates with misfolded proteins in the cytoplasm and mitochondria to provide an environment for protein refolding (33–36). Similar to HSP60, TRiC is a major eukaryotic cytoplasmic chaperonin that is responsible for the correct folding of endogenous client proteins that are prone to misfolding (34,37). The interaction of chaperonin complexes with ADAT3-V144M is consistent with a misfolding and aggregation defect caused by the V144M mutation as suggested above by microscopy and co-IP. We also note that peptides matching ADAT2 were not identified in the purifications of either ADAT3-WT or V144M when purified without co-expression of ADAT2. The lack of copurification of ADAT2 with over-expressed ADAT3 agrees with previous findings that the levels of endogenous ADAT2 are limiting for formation of an ADAT2/3 complex (15).

**Figure 3.**
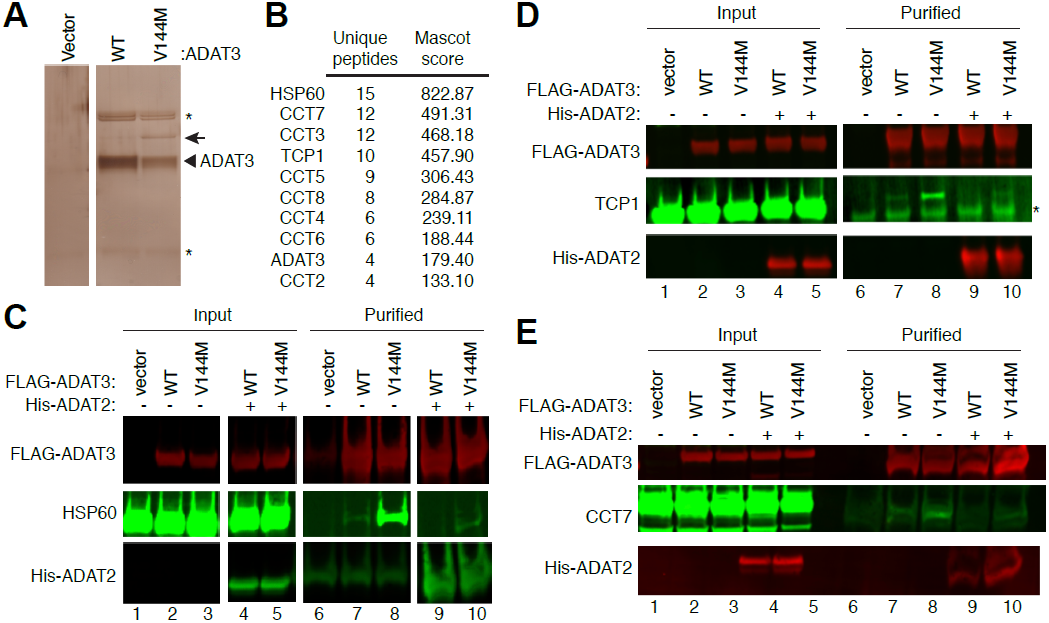
ADAT3-V144M is bound by the HSP60 and TRiC chaperonins. (A) Silver stain of eluted FLAG-affinity purifications from 293T cells expressing FLAG-tag alone (vector), FLAG-ADAT3-WT or FLAG-ADAT3-V144M. Arrowhead represents FLAG-ADAT3 and arrow represents a protein that specifically co-purifies with ADAT3-V144M. (B) Chaperonin proteins identified by LC-MS proteomics specifically in ADAT3-V144M purifications. ADAT3 peptides are included as comparison. (C-E) Immunoblot for the indicated proteins from input (5%) or FLAG-affinity purifications (100%) from 293T cells transfected to express FLAG-ADAT3-WT or V144M without or with His-ADAT2. IP-immunoblots were repeated three times with comparable results. (*) in (A) and (D) represents heavy and light chains of the anti-FLAG antibody used for affinity purification.

To verify and characterize the interaction between HSP60 and ADAT3-V144M, we performed co-IP experiments followed by immunoblotting. For a subset of these assays, we co-expressed His-tagged ADAT2 with either WT or V144M versions of FLAG-ADAT3 to investigate whether ADAT2 influences HSP60 interactions as seen above. While a low level of HSP60 copurified with ADAT3-WT, we detected a significantly increased amount of HSP60 associated with the ADAT3-V144M variant (Figure 3C, HSP60, lanes 7 and 8). Similar to the oligomerization results described above, the interaction between HSP60 and ADAT3-V144M could be greatly suppressed by co-expression with ADAT2 (Figure 3C, compare lanes 8 and 10). The reduction in HSP60 association with ADAT3 by ADAT2 co-expression again demonstrates that assembly of ADAT2 with ADAT3 is likely to prevent aggregation and subsequent targeting by chaperonin complexes. Of note, a similar level of ADAT2 copurified with both ADAT3-WT or V144M (Figure 3C, ADAT2, lanes 9 and 10), further corroborating the results observed above that the V144M mutation does not compromise the interaction between ADAT2 and ADAT3.

Using an analogous co-IP approach, we also found that the TRiC complex subunits TCP1 and CCT7 exhibited significantly increased association with ADAT3-V144M compared to wildtype ADAT3 (Figure 3D and E, lanes 7 and 8). Similar to ADAT3 interaction with HSP60, we also found that co-expression of ADAT2 could suppress the association between the TRiC complex with ADAT3-WT while significantly reducing the amount of TRiC associated with ADAT3-V144M (Figure 3D and E, compare lanes 7 and 8 to 9 and 10). The targeting of either ADAT3-WT or V144M by cellular chaperonin complexes provide evidence that ADAT3-WT is prone to misfolding with the V144M mutation further exacerbating the misfolding and/or aggregation phenotype. Furthermore, these studies reveal a critical role for ADAT2 in facilitating the proper folding of ADAT3 in addition to its role in the nuclear import of assembled ADAT2/3 complexes.

### Purified ADAT2/3 complexes assembled with ADAT3-V144M exhibit defects in adenosine deaminase activity

The aggregation phenotype exhibited by ADAT3-V144M indicates that the V144M mutation is likely to be altering the folding and structure of ADAT3 that could compromise the activity of the ADAT2/3 complex. To probe if the V144M mutation affects ADAT3 activity in wobble inosine formation, we investigated the biochemical properties of the purified ADAT2/3 complexes assembled with either Strep-ADAT3-WT or ADAT3-V144M that were described above (Figure 2B). We analyzed the purified ADAT2/3 complexes for enzymatic activity using an adenosine deaminase assay based upon the separation of digested RNA nucleoside products by thin layer chromatography (TLC) (Figure 4A). For this assay, *in vitro* transcribed tRNA substrates were internally radiolabeled at adenosine residues using [α-^32^P]ATP, incubated with purified ADAT2/3 complexes, digested to nucleoside monophosphates with P1 nuclease and separated by TLC to detect inosine monophosphate (IMP) formation (38,39). As substrates, we used pre-and mature versions of tRNA-Val-AAC since they have been shown to be substrates of ADAT2/3-catalyzed deamination *in vitro* and *in vivo* (15).

**Figure 4.**
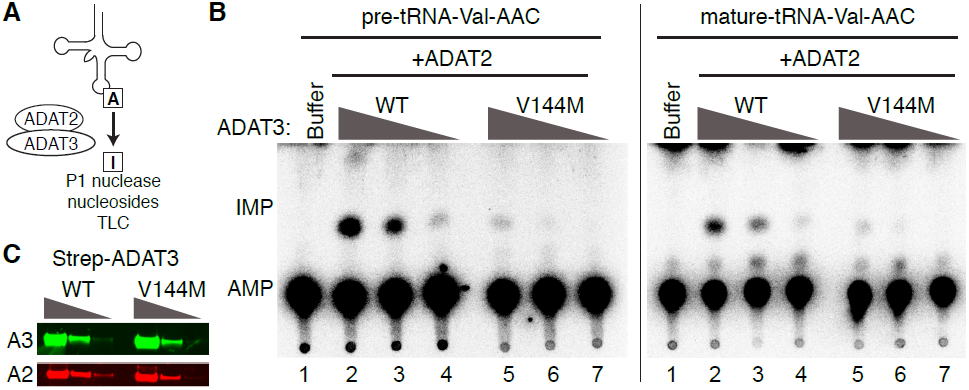
ADAT2/3 complexes assembled with ADAT3-V144M exhibit defects in deaminase activity. (A) Schematic of adenosine deaminase assay for inosine formation using *in vitro* transcribed tRNA-Val-AAC. (B) 5-fold dilution series of purified ADAT2/3 complexes decreasing from 15 nM used for adenosine deaminase activity assays. (C, D) Phosphorimager scans of TLC plates of separated nucleoside products from tRNAs incubated with the indicated buffer or enzymes. The migration of inosine monosphosphate (IMP) and adenosine monophosphate (AMP) are indicated. Adenosine deaminase assays were repeated at least 2x using independently purified ADAT2/3 enzymes with similar results.

Using this assay, we found that purified ADAT2/3 complexes assembled with ADAT3-WT exhibited adenosine deaminase activity on both pre-or mature tRNA-Val-AAC as evidenced by the formation of inosine (Figure 4B and C, IMP, lanes 2-4). In contrast to ADAT2/3-WT complexes, purified ADAT2/3-V144M complexes were significantly diminished in adenosine deaminase activity on either pre- or mature tRNA-Val-AAC (Figure 4B and C, compare lanes 2-4 to lanes 5-7). Based upon the 5-fold dilutions of enzyme, we found that ADAT2/3 complexes assembled with ADAT3-V144M displayed at least a 25-fold decrease in adenosine deaminase activity compared to complexes assembled with ADAT3-WT on either pre- or mature-tRNA-Val-AAC (Figure 4B and C). These findings reveal that while ADAT3-V144M can still associate with ADAT2 in human cells, the variant ADAT2/3-V144M complexes exhibit major defects in adenosine deaminase activity.

### Human patients with homozygous ADAT3-V144M mutations exhibit perturbations in cellular tRNA adenosine deaminase activity

The results thus far provide evidence that ADAT3-V144M exhibits aberrant subcellular localization, increased propensity to aggregate, and defects in adenosine deaminase activity. To examine the molecular effects of the V144M mutation in the human population, we generated lymphoblastoid cell lines (LCLs) from two unrelated human patients harboring homozygous V144M missense mutations in the *ADAT3* gene (referred to as V144M-LCLs generated from patients 1 and 2; P1 and P2; Figure 5A-B). P1 (09DG0640) has been described in detail previously by our group (16,18). Briefly, this is a 6-year old female with severe ID, short stature, microcephaly, strabismus, deafness and history of global developmental delay. She is part of a consanguineous family with three similarly affected siblings, one of whom (09DG00479) is shown in Figure 5C. P2 (11DG1699) is a 24-year old male with severe ID, short stature, microcephaly, strabismus and history of global developmental delay as a child. P2 is part of a consanguineous family with a similarly affected brother (17). LCLs generated from both ID-affected individuals with homozygous V144M mutations were compared to control lymphoblasts from an ethnically matched, healthy, unrelated individual.

**Figure 5.**
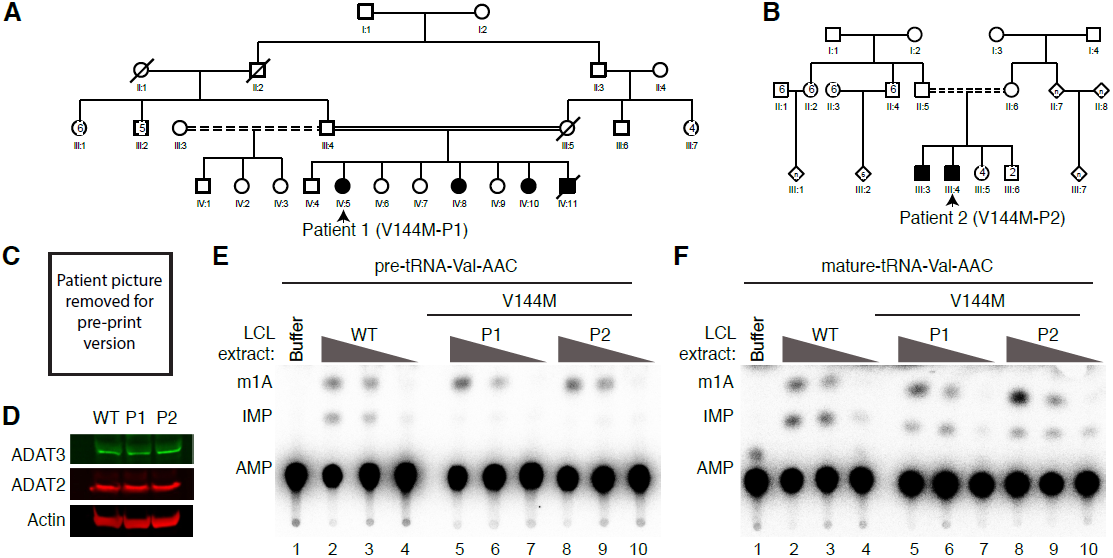
Individuals homozygous for the ADAT3-V144M mutation exhibit defects in adenosine deaminase activity. (A, B) Pedigrees of patients 1 (P1) and 2 (P2) containing homozygous V144M missense mutations in the ADAT3 gene. (C) Frontal and side views of individual 09DG00479 (sibling of P1) showing a prominent forehead, triangular face and strabismus. (D) Immunoblot for the indicated proteins of extracts from LCLs donated from a wildtype individual and patients 1 and 2 harboring homozygous V144M mutations. (E, F) Representative TLC plates from adenosine deaminase activity assays using the indicated tRNA substrates with LCL extracts isolated from the wildtype control individual and persons harboring homozygous V144M mutations. Adenosine deaminase assays were repeated at least 2x with extracts generated from LCLs grown at different times with similar results.

We first examined the levels of ADAT3 protein in the LCLs to determine if the expression or stability of ADAT3 was affected by the V144M mutation. Based upon immunoblotting of whole cell lysates, no major change in the endogenous levels of ADAT3 was detected between WT and V144M-LCLs (Figure 5D, ADAT3). Moreover, we detected no significant change in the levels of the ADAT3 heterodimeric binding subunit, ADAT2, between any of the LCLs (Figure 5D, ADAT2). The comparable steady-state levels of wildtype ADAT3 and the V144M variant suggest the V144M mutation could be affecting ADAT3 function without affecting the levels of protein.

We next employed the TLC-based IMP detection assay described above to investigate whether the V144M mutation affects adenosine deaminase activity in the cells of ID-affected individuals expressing only the ADAT3-V144M variant. Due to the limited amounts of ADAT2/3 found in LCLs combined with the lack of an effective antibody for immunoprecipitation (data not shown), we optimized an *in vitro* adenosine deaminase activity assay using whole cell extracts prepared from the human LCLs described above. This was made possible since previous studies have shown that the ADAT2/3 enzyme complex is the only known cellular activity that catalyzes wobble inosine formation in tRNA (9). Since we found that purified ADAT2/3 complexes exhibit defects in adenosine deaminase activity on pre- and mature tRNA-Val-AAC (Figure 4), we tested whether the LCL extracts also displayed differences in activity on these same substrates.

While no detectable IMP was detected in tRNA substrates pre-incubated with buffer alone, we could readily detect the formation of IMP in both pre- and mature-tRNA-Val-AAC after pre-incubation with whole cell extracts prepared from WT-LCLs (Figure 5E and F, lanes 2-4). In addition to IMP, we also detected a faster-migrating adenosine modification that is consistent with the formation of 1-methyladenosine (Figure 5E, m1A) (38). Since tRNA-Val-AAC has been shown to contain m1A at position 58 in human cells (40), the formation of m1A on the tRNA substrates is likely due to endogenous TRMT6/TRMT61 complexes present in cellular extracts (41). The formation of m1A provides an additional control for normalizing cellular activity since they are catalyzed by two different enzyme complexes.

Agreeing with the activity defect discovered above with purified ADAT3-V144M, we found that V144M-LCL extracts exhibited at least a 25-fold decrease in the generation of inosine in pre-tRNA-Val-AAC (Figure 5E, lanes 5-10). The defect in adenosine deaminase activity for pre-tRNA-Val-AAC was not due to general loss of activity of the V144M-LCL extracts or underloading since the formation of m1A was similar between all three extracts (Figure 5E, m1A). Interestingly, we found that WT- and V144M-LCL extracts displayed a similar level of adenosine deaminase activity on mature tRNA-Val-AAC (Figure 5F). Thus, the V144M mutation appears to affect adenosine deaminase activity on the pre-tRNA-Val-AAC substrate but not processed, mature tRNA-Val-AAC. As an additional control, we compared the level of adenosine deaminase activity in the V144M-LCLs to a completely different LCL line procured from a healthy, wildtype individual of similar age and also detected a similar decrease in tRNA modification activity on pre-tRNA-Val-AAC associated with the V144M-LCL extracts (Supplemental Figure 2). These findings uncover a severe modification defect associated with the ADAT3-V144M variant in human individuals on tRNA-Val-AAC that affects adenosine deaminase activity on the unprocessed form of tRNA greater than the mature form. Importantly, these findings suggest that individuals expressing only the ADAT3-V144M variant are likely to be compromised but not completely abolished in adenosine deaminase activity on wobble inosine-containing tRNA substrates *in vivo*.

### Individuals homozygous for the ADAT3-V144M mutation exhibit decreased wobble inosine modification in some but not all tRNAs

Based upon the adenosine deaminase activity defects exhibited by ADAT2/3-V144M complexes, we next investigated the wobble inosine status of tRNA-Val-AAC isolated from the V144M-LCLs of ID-affected individuals. Since inosine is read as G by reverse transcriptases (42–44), the formation of inosine at the wobble position of tRNA-Val-AAC can be directly detected by sequencing of amplified cDNA obtained by reverse transcription of cellular tRNA. In WT-LCLs, the majority of wobble adenosines in tRNA-Val-AAC were converted to inosine as evidenced by the presence of a predominant “G” peak at position 34 (Figure 6A, Control-WT1). We also compared the level of wobble inosine modification in the WT-LCLs to a completely different LCL line procured from a healthy individual of a similar age and found a similar extent of wobble inosine modification (Figure 6A, Control-WT2). Notably, the level of wobble inosine modification in tRNA-Val-AAC was greatly reduced in both V144M-LCLs with the majority of peak signal at the wobble position being the unmodified “A” (Figure 6A, V144M-P1 and P2). These results provide the first evidence that the ADAT3-V144M mutation and its associated molecular defects have an impact on the levels of tRNA wobble inosine modification *in vivo*.

**Figure 6.**
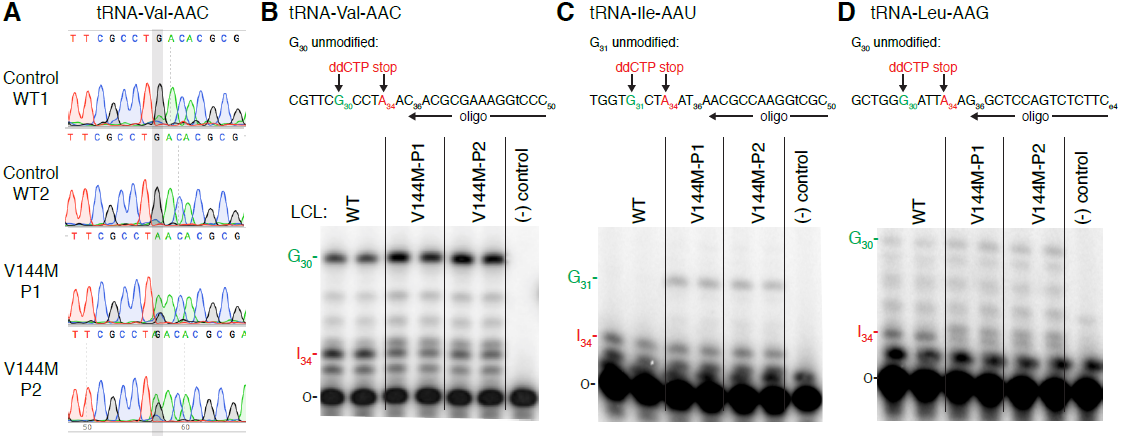
ID-affected individuals expressing only ADAT3-V144M variant exhibit decreased wobble inosine modification in tRNA isoacceptors. (A) Sequencing chromatogram analysis of RT-PCR products amplified from endogenous tRNA-Val-AAC isolated from LCLs of the indicated individuals. The wobble adenosine/inosine position is highlighted. Inosine is read out as G. (B-D) V144M-LCLs exhibit decreased inosine modification in tRNA-Val-AAC, Ile-AAU and Leu-AAG. Primer extension analysis with the indicated oligonucleotide probes against inosine-containing tRNAs in the presence of ddCTP. ‘G _n_’ denotes read-through product indicative of decreased inosine modification at position 34. ‘I_34_’ represents stop position if inosine is present. ‘o’ represents the labeled oligonucleotide used for primer extension.

Based upon the inosine modification defect in tRNA-Val-AAC, we also investigated whether additional tRNAs containing inosine at the wobble position were affected by the ADAT3-V144M mutation. Due to technical challenges in RT-PCR sequencing analysis caused by the diverse number of tRNA isodecoder variants encoded by mammalian genomes, we investigated the modification status of human tRNAs using poisoned primer extension assays with a ddCTP terminator, to distinguish I_34_ (terminated with ddCTP) from A_34_, terminated at the next guanosine (Fig. 6B-D). Using this assay, we observed substantially reduced frequency of I_34_ modification for tRNA-Val-AAC and tRNA-Ile-AAU (Fig. 6B and C), but only a slight defect in modification of tRNA-Leu-AAG (Figure 6D). These results suggest that not all tRNAs are affected to the same extent by the ADAT3-V144M mutation, consistent with the reduced but not abolished adenosine deaminase activity detected in V144M-LCL extracts. Taken altogether, these studies uncover a wobble inosine hypomodification defect for particular tRNAs in the cells of individuals that are homozygous for the ADAT3 V144M mutation, consistent with the multiple perturbations in ADAT3 localization and activity associated with the V144M mutation.

## Discussion

The molecular consequences of the ID-causing ADAT3-V144M mutation have previously been unknown. Surprisingly, the V144M mutation has little to no effect on the steady-state levels of ADAT3 nor on binding to the ADAT2 subunit. However, the ADAT3-V144M mutation significantly alters the subcellular localization properties of ADAT3 and greatly compromises the adenosine deaminase activity of ADAT2/3 complexes assembled with ADAT3-V144M. While the V144M mutation is predicted to be a relatively minor change since valine and methionine represent amino acid residues with hydrophobic side chains of similar molecular size, modeling with the TadA homodimeric complex suggests that the mutation could alter a loop located on the surface of the ADAT3 protein (16). The alteration in ADAT3 structure could then affect the biochemical properties of the assembled ADAT2/3 complex such as substrate binding or adenosine deaminase activity on certain tRNA substrates. Intriguingly, studies with *Trypanosoma brucei* homologs of Tad2p/Tad3p have revealed a role for ADAT3 in substrate tRNA binding and coordination of a single zinc ion (39,45). This would suggest that the ADAT3-V144M mutation may alter the substrate recognition site of the ADAT2/3 complex thereby affecting the binding of or catalysis step on certain tRNA substrates. Indeed, we find that the ADAT3-V144M mutation has differential effects on wobble inosine levels with certain tRNAs exhibiting a substantial decrease in tRNA modification while others display little to no major change. This differential effect could be due to the recognition mechanism of human ADAT2/3, which targets tRNA anticodon loops for inosine modification based upon their structural context rather than simply their sequence alone (46). Further refinement using RNA binding assays and kinetics will provide insight into the specific effect of the V144M mutation on ADAT2/3 enzymatic activity that influence inosine modification levels *in vivo*.

The association of ADAT3-WT with cytoplasmic chaperonins suggests that endogenous ADAT3 could be prone to misfolding and aggregation during translation or after release from the ribosome if not assembled with ADAT2. Consistent with our results, others have found that expression of soluble eukaryotic ADAT3 in *E. coli* requires co-expression of ADAT2 (13,47). The increased tendency of ADAT3-V144M to form cytoplasmic foci suggests that the V144M mutation could further aggravate misfolding and aggregation. Of note, structural studies have shown that methionine differs from other hydrophobic residues in that it can form non-covalent interactions with aromatic-containing residues such as tryptophan, tyrosine or phenylalanine (48,49). In addition, molecular modeling simulation experiments have identified approximately one-third of solved protein structures to contain a methionine-aromatic residue interaction (50). Thus, the substitution of a valine to methionine in the N-terminal extension of ADAT3 could lead to a non-specific interaction with aromatic residues of another ADAT3 protein leading to aggregation. Furthermore, the mutant methionine residue in ADAT3-V144M could form an intramolecular interaction that leads to a different conformation that exacerbates misfolding and aggregation. The misfolding of the ADAT3-V144M protein could reduce the total pool of properly-folded ADAT3 that can interact with ADAT2 leading to decreased levels of active ADAT2/3 complexes. Intriguingly, the perturbed nuclear localization and aggregation into discrete cytoplasmic foci exhibited by the ADAT3-V144M variant is reminiscent of other RNA binding proteins known to misfold and homo-oligomerize in neurological disorders such as TDP-43 and TLS/FUS (51–55).

Due to the intricate dynamics of tRNA processing (56–58), the disruption of nucleocytoplasmic localization by the V144M mutations provides yet another possible contributor to the decreased levels of wobble inosine tRNA modification in ID-affected individuals. The increased propensity of ADAT3-V144M to localize to the nucleus indicates that there could be a relative decrease in the amount of cytoplasmic ADAT3 in ID-affected individuals with homozygous ADAT3-V144M mutations. The cytoplasmic population of ADAT3 assembled with ADAT2 could play a role in modifying tRNAs that have been exported without prior wobble inosine modification by nuclear ADAT2/3. Thus, the disruption of the nucleocytoplasmic ratio by the V144M mutation combined with the activity defect of ADAT2/3-V144M complexes could lead to the reduction in wobble inosine modification levels observed in the tRNAs of individuals homozygous for the ADAT3-V144M mutation. In addition, newly-exported tRNAs lacking inosine could undergo retrograde transport back into the nucleus to be modified by nuclear ADAT2/3 (59,60). Since ADAT2/3-V144M complexes in extract appear to be defective in modifying pre- but not mature tRNA, retrograde transport could play a critical role in providing at least enough tRNA wobble inosine modification for sufficient translation to sustain cell viability.

It is unknown why the ADAT3-V144M mutation causes increased localization to the nucleus. Possible reasons include fortuitous interaction with nuclear import factors, increased interaction with ADAT2 which is responsible for ADAT3 import or loss of interaction with a cytoplasmic retention factor. Investigation of the ADAT3-interactome revealed no obvious candidates that suited any of these scenarios, which is not surprising considering that these factors are likely to be transiently associated. Another possibility is that the increased level of ADAT3 in the nucleus is due to the import of ADAT3 oligomers into the nucleus by ADAT2. Thus, for every ADAT2 imported into the nucleus, there could be more than one ADAT3-V144M subunit translocated alongside.

Based upon the findings, we hypothesize that a certain level of wobble inosine modification in particular tRNAs is necessary for the expression of cellular mRNAs that are critical for proper cellular physiology and human development. Consistent with this prediction, studies in yeast *Schizosaccharomyces pombe* and the plant *Arabodopsis thaliana* have shown that a decrease in tRNA wobble inosine modifications leads to temperature sensitivity, cell cycle arrest and growth retardation (12,14). Moreover, genome wide studies predict numerous highly-expressed genes that are dependent upon ADAT2/3-catalyzed wobble inosine modification for translation (61). Thus, the V144M mutation could alter the cellular proteome in multiple tissues with particularly acute effects in the brain on neural growth and differentiation.

## Materials and Methods

### Human subjects

Evaluation of affected members by a board-certified clinical geneticist included obtaining medical and family histories, clinical examination, neuroimaging and clinical laboratory investigations. After obtaining a written informed con-sent for enrollment in an IRB-approved project (KFSHRC RAC#2070023), venous blood was collected in EDTA and sodium heparin tubes for DNA extraction and establishment of lymphoblastoid cell lines (patients 11DG1699 and 09DG0640, and control 15DG0421), respectively. A separate consent to publish photographs was also obtained.

### Plasmids

The open reading frame for ADAT2 was PCR amplified from cDNA clone RC212395 (Origene) and cloned into pcDNA3.1 (Thermo Fisher) for expression as an untagged protein or as an N-terminal fusion protein with either the 6xHis tag or twin Strep-tag (32). The open reading frame for human ADAT3 was PCR amplified from cDNA plasmid HsCD00326376 (PlasmID Repository, Harvard Medical School) and cloned into either pcDNA3.1-Strep-C, pcDNA3.1-3xFLAG-SBP, pcDNA3.1-N-EGFP, pcDNA3.1-EGFP-C (62). The ADAT3-V144M variant was generated by Gibson mutagenesis using cloning of PCR fragments and verified by Sanger sequencing.

### Cell culture

HeLa S3 human cervical carcinoma and 293T human embryonic kidney cell lines were cultured in DulbeccO’s Minimal Essential Medium (DMEM) containing 10% fetal bovine serum, 2 mM L-alanyl-L-glutamine (GlutaMax, Gibco) and 1% Penicillin/Streptomycin. Human lymphoblastoid cell lines were cultured in Roswell Park Memorial Institute (RPMI) 1640 Medium containing 15% fetal bovine serum, 2 mM L-alanyl-L-glutamine (GlutaMax, Gibco) and 1% Penicillin/Streptomycin. The additional wildtype control LCL line was obtained from Coriell Institute for Medical Research #GM22647.

### Microscopy

Hela cells were plated at 2.5 ×10^5^cells on a 6-well plate. Cells were transfected 1 day after plating with a total of 2.5 μg of DNA using Lipofectamine 3000. Cells were imaged 48 hours post-transfection on an EVOS fluorescence microscopy imaging system (ThermoFisher). For DNA staining, cells were washed twice with PBS and then incubated for 30 minutes at 37 degrees with PBS containing 10% FBS and 1 μM of Hoechst and then imaged. For cytoplasmic foci quantification, 5 images were taken of each well and the number of GFP-positive cells along with the number of cells containing greater than 3 foci were counted in each of the 5 frames. The experiment was performed 3x on N-terminal GFP-tagged ADAT3 with a minimum of 580 cells counted per experiment and independently verified by blinded analysis. For C-terminal GFP-tagged ADAT3, the experiment was performed 2x with independent verification. For quantification of nuclear ADAT3, each of the three N-terminal experiments were quantified using the same minimum of 580 cells counted per experiment.

### Protein purification and analysis

Transient transfection and cellular extract production were performed as previously described (63). In brief, 2.5×10^6^ 293T HEK cells were transiently transfected by calcium phosphate DNA precipitation with 10-20 μg of plasmid DNA followed by preparation of lysate by hypotonic freeze-thaw lysis 48 hours post-transfection. For anti-FLAG purification, whole cell extract from transiently transfected cells cell lines (1 mg of total protein) was rotated with 20 μL of Anti-DYKDDDDK Magnetic Beads (Takara BioUSA, Clontech or Syd labs) for 2 h at 4° C in lysis buffer (20 mM HEPES at pH 7.9, 2 mM MgCl2, 0.2 mM EGTA, 10% glycerol, 1 mM DTT, 0.1 mM PMSF, 0.1% NP-40) with 200 mM NaCl. Resin was washed three times using the same buffer followed by RNA extraction or protein analysis. Strep-tagged proteins were purified using MagSTREP “type3” XT beads, 5% suspension (IBA Lifesciences) under similar conditions as with anti-FLAG purifications and eluted with desthiobiotin.

Protein identification was performed by the URMC Mass Spectrometry Resource Laboratory. Briefly, protein samples were reduced, alkylated and digested in solution with trypsin followed by purification and desalting on an analytical C18 column tip. Peptide samples were analyzed by HPLC chromatography coupled with electrospray ionization on a Q Exactive Plus Hybrid Quadrupole-Orbitrap mass spectrometer (Thermo Fisher). Protein identification through tandem mass spectra correlation was performed using SEQUEST and Mascot.

Cellular extracts and purified protein samples were fractionated on NuPAGE Bis-Tris polyacrylamide gels (Thermo Scientific) followed by transfer to Immobilon FL PVDF membrane (Millipore) for immunoblotting. For analysis of LCL extracts, 5×10^6^ lympoblast cells were harvested and proteins were extracted using radioisotope immunoprecipitation assay (RIPA) buffer (50 mM TrisHCl, pH 7.5, 1% NP-40, 0.5% sodium deoxycholate, 0.1% SDS, 150 mM NaCl, 2mM EDTA). Antibodies were against the following proteins : FLAG epitope tag (L00018, Sigma), 6xHis tag (MA1-21315, Thermo Fisher), GFP (sc-9996, Santa Cruz Biotechnology), Strep-tag II-tag (NC9261069, Thermo Fisher), ADAT3 (Abcam, ab192987), ADAT3 (H00113179-B01P, Abnova), ADAT2 (ab135429, Abcam), HSP60 (A302-845A, Bethyl Labs), CCT1 (sc-53454, Santa Cruz Biotechnologies), CCT7 (A304-730A-M, Bethyl Labs). Primary antibodies were detected using IRDye 800CW Goat anti-Mouse IgG (925-32210, Thermofisher) or Rabbit (SA5-35571, Thermofisher) or Rat (925-32219, LI-COR Biosciences), or IRDye 680RD Goat anti-Mouse IgG (926-68070, LI-COR Biosciences) or Rabbit (925-68071). Immunoblots were scanned using direct infrared fluorescence via the Odyssey System (LI-COR Biosciences).

### Adenosine deaminase assays

Internally-radiolabeled tRNA substrates were prepared by T7 *in vitro* transcription of DNA templates generated by PCR amplification. Oligonucleotides containing the T7 promoter upstream of tRNA sequences were PCR amplified using Herculase II DNA polymerase or Taq DNA Polymerase (New England Biolabs) followed by agarose gel purification of PCR amplification products. *In vitro* transcription was performed using Optizyme T7 RNA polymerase (Fisher Scientific) with 10 mM each of UTP, CTP and GTP, 1mM of ATP and 250 μCi of [α-^32^P]-ATP (800Ci/mmol, 10mCi/ml). *In vitro* transcription reactions were incubated at 37° C for 2 hours followed by DNase treatment and purification using RNA Clean and Concentrator columns (Zymo Research). Full-length tRNA transcripts were verified on a 15% Polyacrylmide-urea gel stained with SYBR Gold nucleic acid stain (Thermo Fisher). Before conducting enzymatic assays, all tRNA substrates were refolded by thermal denaturation at 95°C for 2 minutes in buffer containing a final concentration 5 mM TRIS ph7.5 and 0.16 mM EDTA, quick chilling on ice for 2 min and refolding at 37°C in the presence of Hepes pH 7.5, MgCl2, and NaCl.

For adenosine deaminase assays, ∼30 ng of refolded tRNA substrate was incubated with titrations of enzyme starting at the highest concentration of ∼15 nM of Strep-purified ADAT3. Serial dilutions occurring in 1/5 increments then were carried out in the presence of 12.5 μg/mL BSA. For enzymatic reactions from extracts, the highest concentration of protein started at ∼20 ug of total protein in the reaction. Dilutions occurring in 1/5 increments were then carried out. Reactions were incubated at 37°C for 60 minutes and RNA was purified using RNA Clean and Concentrator columns. The tRNA was eluted in 20 μL of water and 10 μL was subjected to nuclease P1 digestion overnight in total volume of 13 μL and 0.125 units of P1 in 250 mM Ammonium Acetate pH 5.35. Half of the P1-nuclease treated samples were spotted on a POLYGRAM^®^ polyester Cellulose MN 300 plates (Macherey Nagel) run in solvent B (0.1M sodium phosphate buffer pH 6.8:NH4 sulfate:n-propanol (100:60L2 [vLw:v]). Phosphorimaging was conducted on a Bio-Rad Personal Molecular Imager followed by analysis using NIH ImageJ software.

### RNA analysis

RNA extraction was performed on 10 × 10^6^ human lyphoblastoid cell lines using Trizol LS reagent (Thermo Fisher). For RT-PCR, total RNA (∼1.25 μg) was reverse transcribed for tRNA-Val-AAC using Superscript IV enzyme followed by the QIAquick PCR purification kit. cDNA was then PCR amplified using Herculase II DNA polymerase (Agilent Genomics). The PCR product was gel extracted and analyzed by Sanger sequencing (ACGT, Inc). The following primers were used:

Val RT primer: TGTTTCCGCCTGGTTTTG

Val PCR primer F: GAACTAAGCTTGTTCAGAGTTCTACAGTCCGGACTACAAAGACCATGACGGTGATTATAAAGATCATGACATGTTTCCGTAGTGTAGTGGTTATCAC Val PCR primer R: CACT TGTTTCCGCCTGGTTTTGATCCAGGGACC

For primer extension assays to monitor inosine modification status, oligonucleotides were 5’ end labeled and purified as previously described (64). In a 5 μL annealing reaction, 0.25 – 1 pmol of labeled primers were annealed to 0.6 μg of bulk RNA by incubation for 3 min at 95°C followed by slow cooling and incubation for 30 min at 50-55°C. The annealing product was then extended using 64 U Superscript III (Invitrogen) in a 10 μL reaction containing 1X First Strand buffer, 2 mM ddCTP, 0.5 mM of each of the other dNTPs and 10 mM MgCl_2_ at 50 - 55°C for 1 h. Reactions were stopped by addition of 2 X RNA loading dye containing 98% formamide, 10 mM EDTA, 1 mg/mL bromophenol blue, and 1 mg/mL xylene cyanol, resolved on a 7M urea - 15% polyacrylamide gel, and the dried gel was imaged on a Typhoon phosphorimager and quantified as described (65). The primer sequences are as noted:

Human tA(IGC) [51-36] CGCTaCCTCTCGCATG

Human tV(AAC) [50-36] GGGaCCTTTCGCGTG

Human tIle(AAT)[50-36] GCGaCCTTGGCGTTA

## Supplementary Materials

Supplemental Figures 1-2 and Supplemental Table 1 are provided.

## Acknowledgements

We thank Mais Hashem for her assistance as a clinical research coordinator; Tarfa Alshiddi, Rana Alomar and Eman Alobaid from the tissue culture core facility at KFSHRC; Joshua Dewe, Morgan Thomalla and Michael Haft for preliminary studies; Kyle Swovick, Chen Chen, and Ben Phelan for foci quantification; Kevin Welle and the URMC Mass Spectrometry Resource Lab for proteomics; and the Ghaemmaghami Lab for microscopy resources and discussion. We also thank the participating human subjects for donating their blood samples for research.

---

**Supplemental Figure 1.**
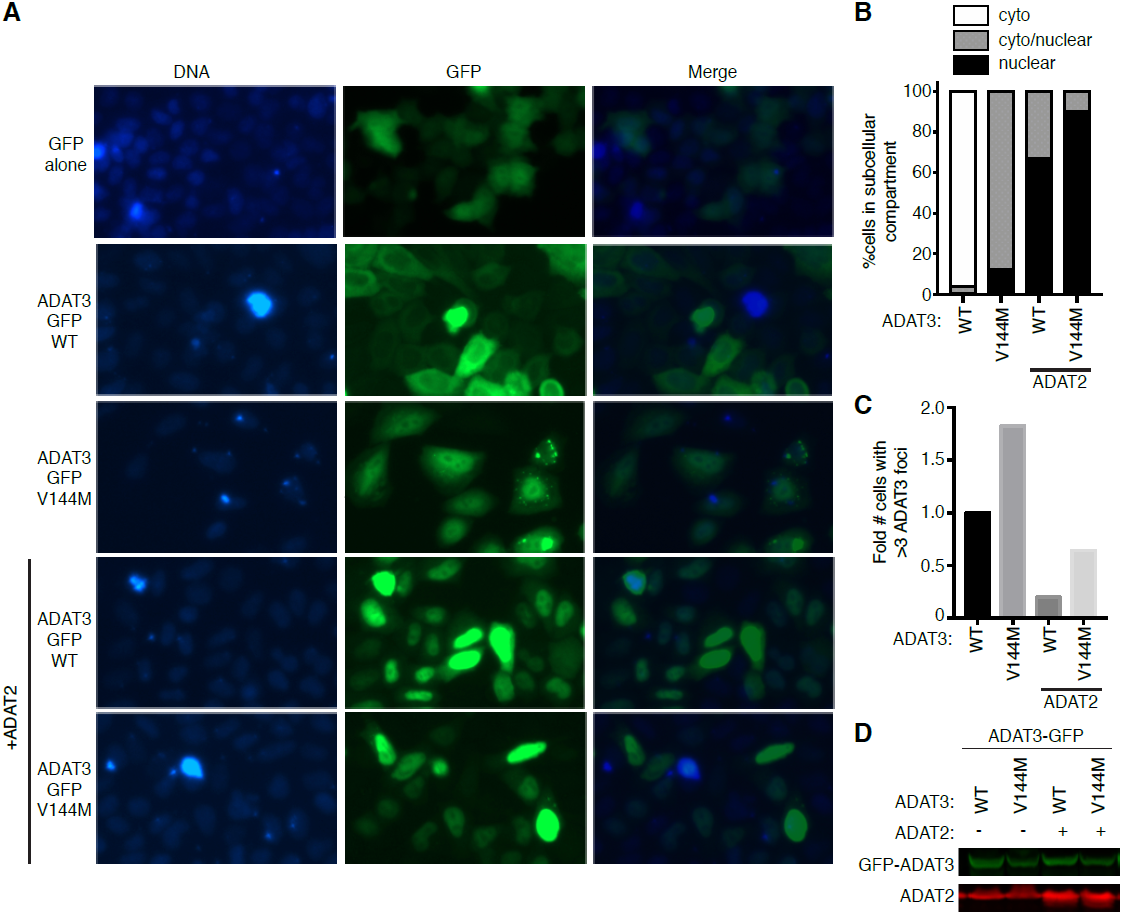
ADAT3-V144M displays aberrant nucleocytoplasmic localization and increased susceptibility to form cytoplasmic aggregates also when tagged on the C-terminus with GFP (ADAT3-GFP). (A) Fluorescence microscopy images of GFP alone, ADAT3-WT and V144M GFP-tagged expressed in HeLa cervical carcinoma cells. Nuclear DNA was stained with Hoechst with merge on right column. (B) Fraction of cells exhibiting ADAT3-GFP that was either primarily cytoplasmic, similarly distributed between the cytoplasmic and nucleus, or primarily nuclear. (C) Fold number of cells that exhibited greater than three cytoplasmic foci of GFP-ADAT3. For (B) and (C) a minimum of 615 cells counted per experiment. (D) Immunoblot of ADAT3-GFP expression without or with ADAT2 co-expression.

**Supplemental Figure 2.**
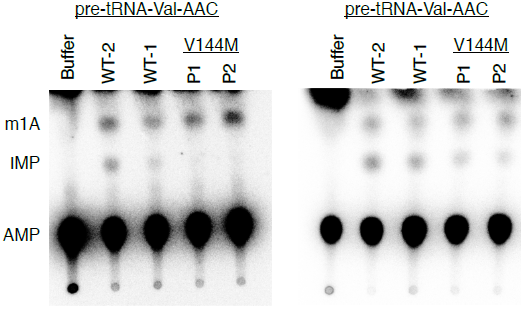
Individuals homozygous for the ADAT3-V144M mutation exhibit defects in adenosine deaminase activity relative to wildtype (WT) control LCLs. TLC plates from adenosine deaminase activity assays using the indicated tRNA substrates with LCL extracts isolated from an additional, unrelated wildtype control (WT-2) individual, the previously-described wildtype control individual (WT-1) and patients harboring homozygous V144M mutations (P1 and P2).

